# Rapid development of an inactivated vaccine for SARS-CoV-2

**DOI:** 10.1101/2020.04.17.046375

**Authors:** Qiang Gao, Linlin Bao, Haiyan Mao, Lin Wang, Kangwei Xu, Minnan Yang, Yajing Li, Ling Zhu, Nan Wang, Zhe Lv, Hong Gao, Xiaoqin Ge, Biao Kan, Yaling Hu, Jiangning Liu, Fang Cai, Deyu Jiang, Yanhui Yin, Chengfeng Qin, Jing Li, Xuejie Gong, Xiuyu Lou, Wen Shi, Dongdong Wu, Hengming Zhang, Lang Zhu, Wei Deng, Yurong Li, Jinxing Lu, Changgui Li, Xiangxi Wang, Weidong Yin, Yanjun Zhang, Chuan Qin

## Abstract

The COVID-19 pandemic caused by SARS-CoV-2 has brought about an unprecedented crisis, taking a heavy toll on human health, lives as well as the global economy. There are no SARS-CoV-2-specific treatments or vaccines available due to the novelty of this virus. Hence, rapid development of effective vaccines against SARS-CoV-2 is urgently needed. Here we developed a pilot-scale production of a purified inactivated SARS-CoV-2 virus vaccine candidate (PiCoVacc), which induced SARS-CoV-2-specific neutralizing antibodies in mice, rats and non-human primates. These antibodies potently neutralized 10 representative SARS-CoV-2 strains, indicative of a possible broader neutralizing ability against SARS-CoV-2 strains circulating worldwide. Immunization with two different doses (3μg or 6 μg per dose) provided partial or complete protection in macaques against SARS-CoV-2 challenge, respectively, without any antibody-dependent enhancement of infection. Systematic evaluation of PiCoVacc via monitoring clinical signs, hematological and biochemical index, and histophathological analysis in macaques suggests that it is safe. These data support the rapid clinical development of SARS-CoV-2 vaccines for humans.

**One Sentence Summary:** A purified inactivated SARS-CoV-2 virus vaccine candidate (PiCoVacc) confers complete protection in non-human primates against SARS-CoV-2 strains circulating worldwide by eliciting potent humoral responses devoid of immunopathology

## Main Text

The World Health Organization declared the outbreak of coronavirus disease in 2019 (COVID-19) to be a Public Health Emergency of International Concern on 30 January 2020, and a pandemic on 11 March 2020. It is reported that ~80% of COVID-19 patients have mild-to-moderate symptoms, while ~20% develop serious manifestations such as severe pneumonia, acute respiratory distress syndrome (ARDS), sepsis and even death (*1*). The number of COVID-19 cases has increased at a staggering rate globally. As of 10 April, the total confirmed cases have reached 1,623, 173 and the death toll has risen to 97,236. Severe acute respiratory syndrome coronavirus 2 (SARS-CoV-2), the causative agent of the ongoing pandemic, belongs to the genus *Betacoronavirus* (β-CoV) of the family *Coronavirdae* (*2*). SARS-CoV-2 along with the severe acute respiratory syndrome coronavirus (SARS-CoV) and the Middle Eastern respiratory syndrome-related coronavirus (MERS-CoV), constitute the three most life-threatening species among all human coronaviruses. SARS-CoV-2 harbors a linear single-stranded positive sense RNA genome, encoding 4 structural proteins [spike (S), envelope (E), membrane (M), and nucleocapsid (N)] of which S is a major protective antigen that elicits highly potent neutralizing antibodies (NAbs), 16 non-structural proteins (nsp1-nsp16) and several accessory proteins (*3*). No specific antiviral drugs or vaccines against the newly emerged SARS-CoV-2 are currently available. Therefore, urgency in the development of vaccines is of vital importance to curb the pandemic and prevent new viral outbreaks.

Multiple SARS-CoV-2 vaccine types, such as DNA-, RNA-based formulations, recombinant-subunits containing viral epitopes, adenovirus-based vectors and purified inactivated virus are under development (*4–6*). Purified inactivated viruses have been traditionally used for vaccine development and such vaccines have been found to be safe and effective for the prevention of diseases caused by viruses like influenza virus and poliovirus (*7, 8*). To develop preclinical *in vitro* neutralization and challenge models for a candidate SARS-CoV-2 vaccine, we isolated SARS-CoV-2 strains from the bronchoalveolar lavage fluid (BALF) samples of 11 hospitalized patients (including 5 ICU patients), among which 4 are from China, 4 from Italy, 1 from Switzerland, 1 from UK and 1 from Spain (table. S1). These patients were infected with SARS-CoV-2 during the most recent outbreak. The 11 samples contained SARS-CoV-2 strains are widely scattered on the phylogenic tree constructed from all available sequences, representing, to some extent, the circulating populations (Fig. 1A and fig. S1). We chose strain CN2 for purified inactivated SARS-CoV-2 virus vaccine development (PiCoVacc) and other 10 strains (termed as CN1, CN3-CN5 and OS1-OS6) as preclinical challenge strains. A number of strains amongst these, including CN1 and OS1, which are closely related to 2019-nCoV-BetaCoV /Wuhan/WIV04/2019 and EPI_ISL_412973, respectively, have been reported to cause severe clinical symptoms, including respiratory failure, requiring mechanical ventilation (*9, 10*).

**Fig. 1.**
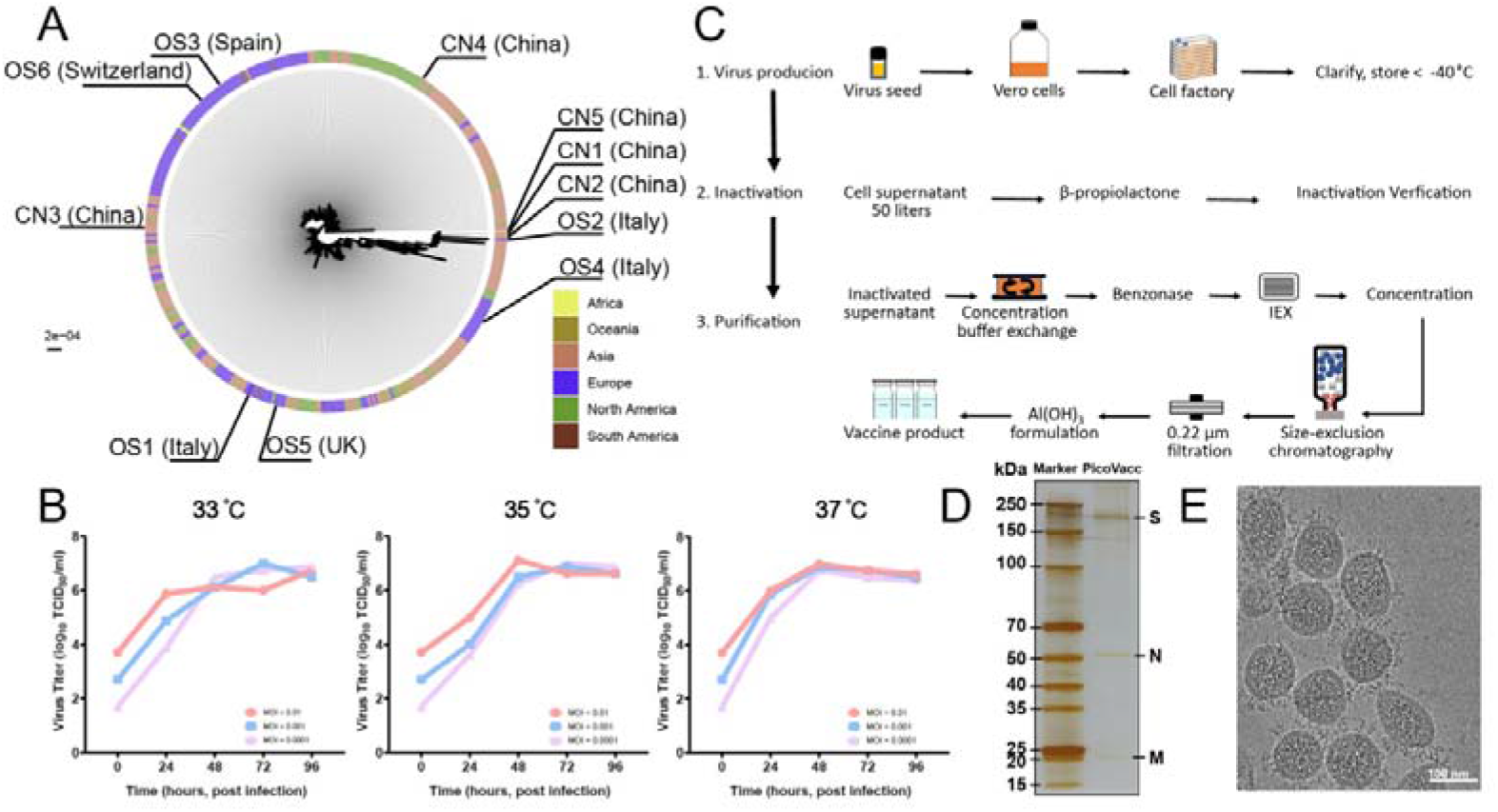
Characterization of PiCoVacc. (**A**) SARS-CoV-2 maximum likelihood phylogenetic tree. The SARS-CoV-2 isolates used in this study are depicted with black lines and labeled. The continents where the virus strains were from are colored differently. (**B**) Growth kinetics of PiCoVacc (CN2) P5 stock in Vero cells. (**C**) Flowchart of PiCoVacc preparation. (**D**) Protein composition and purity evaluation of PiCoVacc by NuPAGE 4-12% Bis-Tris Gel. (**E**) Representative electron micrograph of PiCoVacc.

To obtain a viral stock adapted for efficient growth in Vero cells for PiCoVacc production, the CN3 strain was firstly plaque purified and passaged once in Vero cells to generate the P1 stock. After this another four passages were performed to generate the P2-P5 stocks. Growth kinetics analysis of the P5 stock in Vero cells showed that this stock replicated efficiently and reached a peak titer of 6-7 log_10_ TCID_50_/ml by 3 or 4 days post infection (dpi) at multiplicities of infection (MOI) of 0.0001-0.01 at temperatures between 33°C-37°C (fig. 1B). To evaluate the genetic stability of PiCoVacc, 5 more passages were performed to obtain the P10 stock, whole genome of which, together with those of the P1, P3 and P5 stocks were sequenced. Compared to P1, only two amino acid substitutions, Ala → Asp at E residue 32 (E-A32D) and Thr → Ile at nsp10 residue 49 (nsp10-T32I), occurred in P5 and P10 stocks (table. S2), suggesting that PiCoVacc CN2 strain possesses excellent genetic stability without any 5 mutations that might potentially alter the NAb epitopes. To produce pilot scale PiCoVacc for animal studies, the virus was propagated in a 50-liter culture of Vero cells using cell factory and inactivated by using β-propiolactone (Fig. 1C). The virus was purified using depth filtration and two optimized steps of chromatography, yielding a highly pure preparation of PiCoVacc (Fig. 1D). Additionally, cryo-electron microscopy (cryo-EM) analysis showed intact oval-shaped particles with diameters of 90-150 nm, which are embellished with crown-like spikes, representing a prefusion state of the virus (Fig. 1E).

To assess the immunogenicity of PiCoVacc, groups of BALB/c mice (n=10) were injected at day 0 and day 7 with various doses of PiCoVacc mixed with alum adjuvant (0, 1.5 or 3 or 6 μg per dose, 0 μg in physiological saline as the sham group). No inflammation or other adverse effects were observed. Spike-, receptor binding domain (RBD)-, and N-specific antibody responses were evaluated by enzyme-linked immunosorbent assays (ELISAs) at weeks 1-6 after initial immunization (fig. S2). SARS-CoV-2 S-and RBD-specific immunoglobulin G (Ig G) developed quickly in the serum of vaccinated mice and peaked at the titer of 819,200 (>200 μg/ml) and 409,600 (>100 μg/ml), respectively, at week 6 (Fig. 2A). RBD-specific Ig G accounts for half of the S induced antibody responses, suggesting RBD is the dominant immunogen, which closely matches the serological profile of the blood of recovered COVID-19 patients (Fig. 2A and 2B) (*11*). Surprisingly, the amount of N-specific Ig G induced is ~30-fold lower than the antibodies targeting S or RBD in immunized mice. Interestingly, previous studies have shown that the N-specific Ig G is largely abundant in the serum of COVID-19 patients and serves as one of the clinical diagnostic markers (*11*). It’s worthy to note that PiCoVacc could elicit much higher S-specific antibody titers than those of the serum from the recovered COVID-19 patients. This observation coupled with the fact that the antibodies targeting N of SARS-CoV-2 do not provide protective immunity against the infection (*12*), suggest that PiCoVacc is capable of eliciting more effective antibody responses (Fig. 2A and 2B). Next, we measured SARS-CoV-2-specific neutralizing antibodies over a period of time using microneutralization assays (MN50). Similar to S-specific Ig G responses, the neutralizing antibody titer against the CN1 strain emerged at week 1 (12 for high dose immunization), surged after the week 2 booster and reached up to around 1,500 for low and medium doses and 3,000 for high dose at week 7, respectively (Fig. 2A). In contrast, the sham group did not develop detectable SARS-CoV-2-specific antibody responses (Fig. 2A and 2B). In addition, immunogenic evaluations of PiCoVacc in Wistar rats with the same immunization strategy yielded similar results – the maximum neutralizing titers reached 2,048-4,096 at week 7 (Fig. 2C). To investigate the spectrum of neutralizing activities elicited by PiCoVacc, we conducted neutralization assays against the other 9 isolated SARS-CoV-2 strains using mouse and rat serums collected 3 weeks post vaccination. Neutralizing titers against these strains demonstrate that PiCoVacc is capable of eliciting antibodies that possibly exhibit potent neutralization activities against SARS-Cov-2 strains circulating worldwide (Fig. 2D and 2E).

**Fig. 2.**
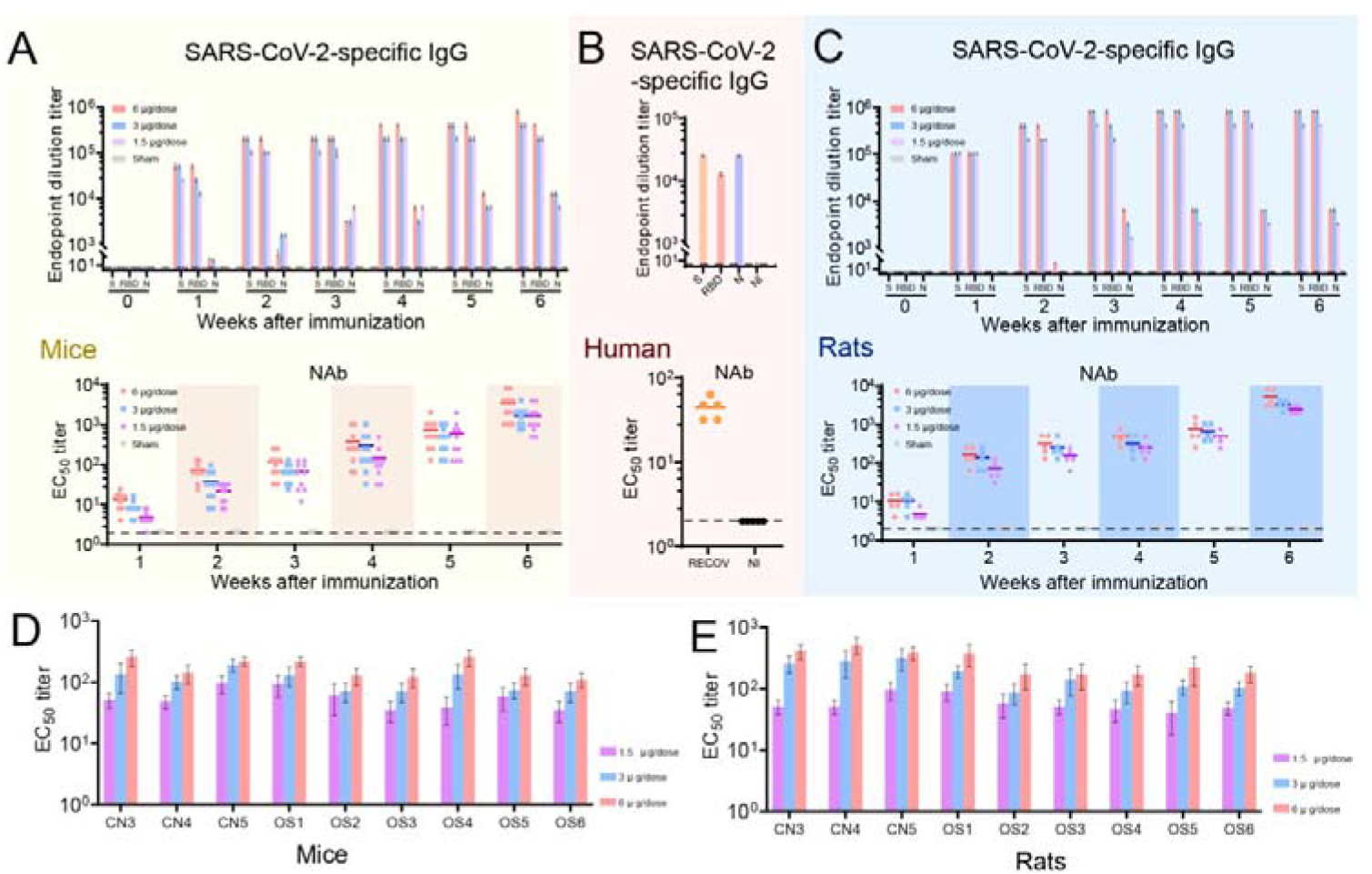
PiCoVacc immunization elicits neutralizing antibody response against 10 representative SARS-CoV-2 isolates. BALB/c mice and Wistar rats were immunized with various doses of PiCoVacc or control (adjuvant only as the sham group) (n=10). Serums from recovered COVID19 patients (RECOV) and non-infected people (NI) act as positive and negative controls, respectively. The antibody responses were analyzed in mice (**A**), humans (**B**) and rats (**C**). Top: SARS-CoV-2-specific IgG response measured by ELISA; bottom: neutralizing antibody titer determined by microneutralization assay. The spectrum of neutralizing activities elicited by PiCoVacc was investigated in mice (**D**) and rats (**E**). Neutralization assays against the other 9 isolated SARS-CoV-2 strains were performed using mouse and rat serums collected 3 weeks post vaccination. Points represent individual animals and humans; dotted lines indicate the limit of detection; horizontal lines indicate the geometric mean titer (GMT) of EC_50_ for each group.

We next evaluated the immunogenicity and protective efficacy of PiCoVacc in rhesus macaques *(Macaca mulatta),* a non-human primate species that shows a COVID-19-like disease caused by SARS-CoV-2 infection *(13).* Macaques were immunized three times via the intramuscular route with medium (3 μg per dose) or high doses (6 μg per dose) of PiCoVacc at day 0, 7 and 14 (n=4). S-specific IgG and NAb were induced at week 2 and rose to ~12,800 and ~50, respectively at week 3 (before virus challenge) in both vaccinated groups, whose titers are similar to those of serum from the recovered COVID-19 patients (Fig. 3A and 3B). Unexpectedly, NAb titer (61) in medium dose immunized group was ~20% more than that observed (50) in high dose vaccinated group at week 3, possibly due to individual differences in the ability of one animal in medium dose group in eliciting ~ 10-fold higher titer when compared to the other 3 (Fig. 3B). Excluding this exception, NAb titer in medium dose group would drop down to 34, ~40% lower than that in high dose group. Subsequently, we conducted a challenge study by a direct inoculation of 10^6^ TCID_50_ of SARS-CoV-2 CN1 into the animal lung through intratracheal route at day 22 (one day after the third immunization) in vaccinated and control macaques to verify the protective efficacy. Expectedly, all control (sham and placebo) macaques showed excessive copies (10^4^-10^6^/ml) of viral genomic RNA in pharynx, crissum and lung by day 3-7 post-inoculation (dpi) and severe interstitial pneumonia (Fig. 3C–3F). By contrast, all vaccinated macaques were largely protected against SARS-CoV-2 infection with very small histopathological changes in lung, which probably were caused by a direct inoculation of 10^6^ TCID_50_ of virus into the lung through intratracheal route, that needed longer time (more than one week) to recover completely (Fig. 3F). Viral loads decreased significantly in all vaccinated macaques, but increased slightly in control animals from day 3-7 after infection (Fig. 3C–3E). All four macaques that received the high dose, had no detectable viral loads in pharynx, crissum and lung at day 7 after infection. In medium dose immunized group, we indeed partially detected the viral blip from pharyngeal (3/4), anal (2/4) and pulmonary (1/4) specimens at day 7 after infection, whilst viral loads presented a ~95% reduction when compared to the sham groups (Fig. 3C–3E). Interestingly, NAb titer in vaccinated groups decreased by ~30% by 3 days post infection to neutralize viruses, then rapidly increased from day 5-7 after infection to maintain its potent neutralization efficacy. In comparison with high dose vaccination group (titer of ~145), higher NAb titers observed in medium dose vaccinated group at day 7 after infection (~400 for 4 macaques; ~300 for 3 macaques, if the one outlier is discarded) might have resulted from relatively low level of viral replication, suggesting a requirement of longer time for complete viral clearance. No antibody-dependent enhancement of infection (ADE) was observed for any vaccinated macaques despite the observation that relatively low NAb titer existed within the medium dose group before infection, offering partial protection.

**Fig. 3.**
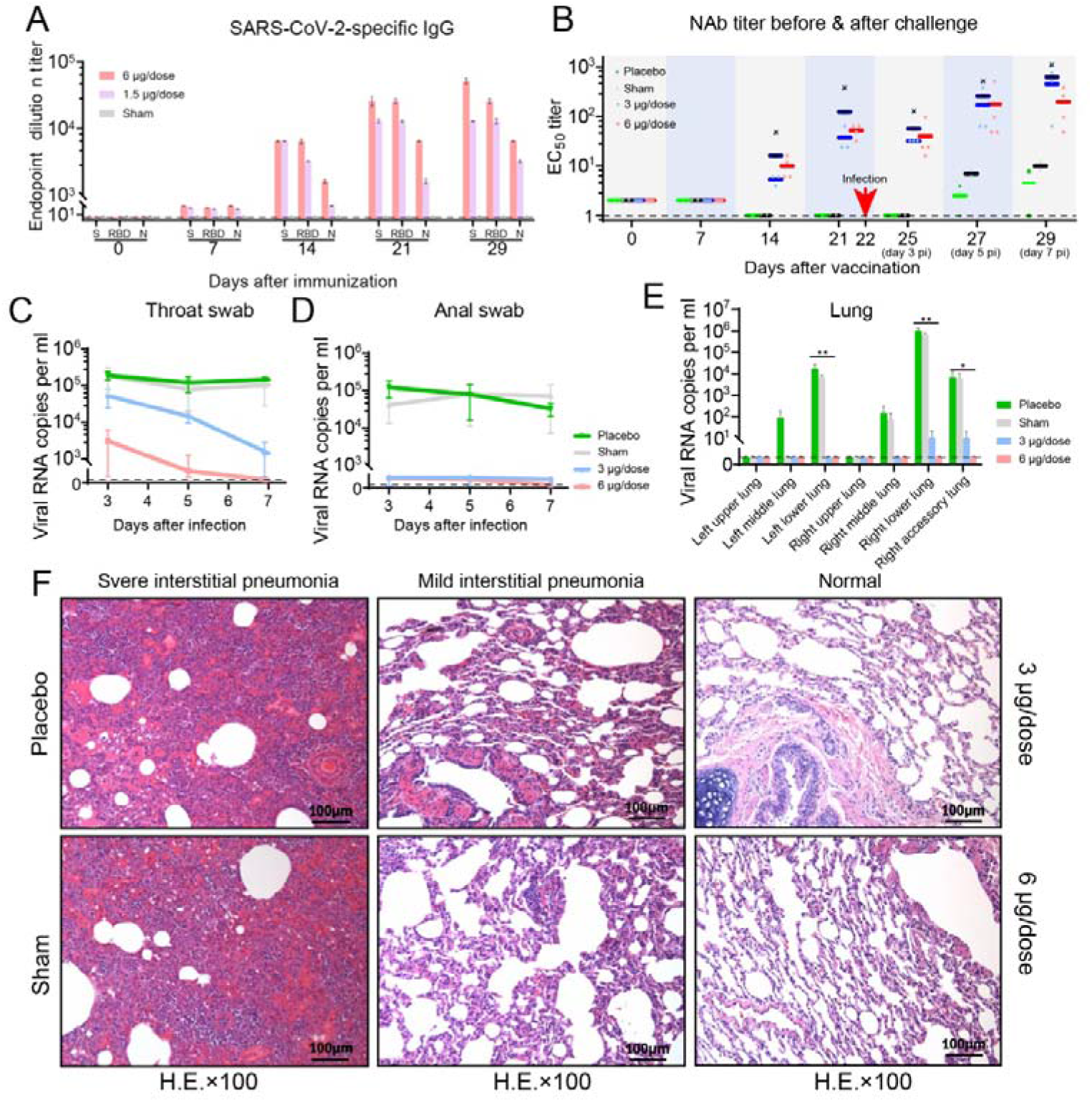
Immunogenicity and protective efficacy of PiCoVacc in nonhuman primates. Macaques were immunized three times through the intramuscular route with various doses of PiCoVacc or adjuvant only (sham) or placebo (n=4). SARS-CoV-2-specific IgG response (**A**) and neutralizing antibody titer (**B**) were measured. Points represent individual macaques; dotted lines indicate the limit of detection; horizontal lines indicate the geometric mean titer (GMT) of EC_50_ for each group. Protective efficacy of PiCoVacc against SARS-CoV-2 challenge at week 3 after immunization was evaluated in macaques (**C-F**). Viral loads of throat (**C**) and anal (**D**) swab specimens collected from the inoculated macaques at day 3, 5 and 7 pi were monitored. Viral loads in various lobe of lung tissue from all the inoculated macaques at day 7 post infection were measured (**E**). RNA was extracted and viral load was determined by qRT-PCR. All data are presented as mean ± SEM. Histopathological examinations (**F**) in lungs from all the inoculated macaques at day 7 post infection. Lung tissue was collected and stained with hematoxylin and eosin.

Previous experiences with the development of SARS and MERS vaccine candidates had raised concerns about pulmonary immunopathology, either directly caused by a type 2 helper T-cell (Th2) response or as a result of ADE (*4, 14, 15*). Although T-cell responses elicited by many vaccines have been demonstrated to be crucial for acute viral clearance, protection from subsequent coronavirus infections is largely mediated by humoral immunity (*16, 17*). The “cytokine storm” induced by excessive T-cell responses have been actually shown to accentuate the pathogenesis of COVID19 (*18, 19*). Therefore, T-cell responses elicited by SARS-CoV-2 vaccine have to be well controlled in order to avoid immunopathology. In context with this, we systematically evaluated safety of PiCoVacc in macaques by recording a number of clinical observations and biological indices. Two groups of macaques (n=10) were immunized by intramuscular injection with low (1.5 μg) or high doses (6 μg) and another two groups of macaques (n=10) were immunized with adjuvant (sham) and physiological saline (placebo) for three times at day 0, 7 and 14. Neither fever nor weight loss was observed in any macaque after the immunization of PiCoVacc, and the appetite and mental state of all animals remained normal (Fig. S3). Hematological and biochemical analysis, including biochemical blood test, lymphocyte subset percent (CD3^+^, CD4^+^ and CD8^+^) and key cytokines (TNF-α, IFN-γ, IL-2, IL-4, IL-5 and IL-6) showed no notable changes in vaccinated groups when compared to the sham and placebo groups (Fig. 4A–4B and fig. S4–S5). In addition, histopathological evaluations of various organs, including lung, heart, spleen, liver, kidney and brain, from the 4 groups at day 29 demonstrated that PiCoVacc did not cause any notable pathology in macaques (Fig. 4C and fig. S6).

**Fig. 4.**
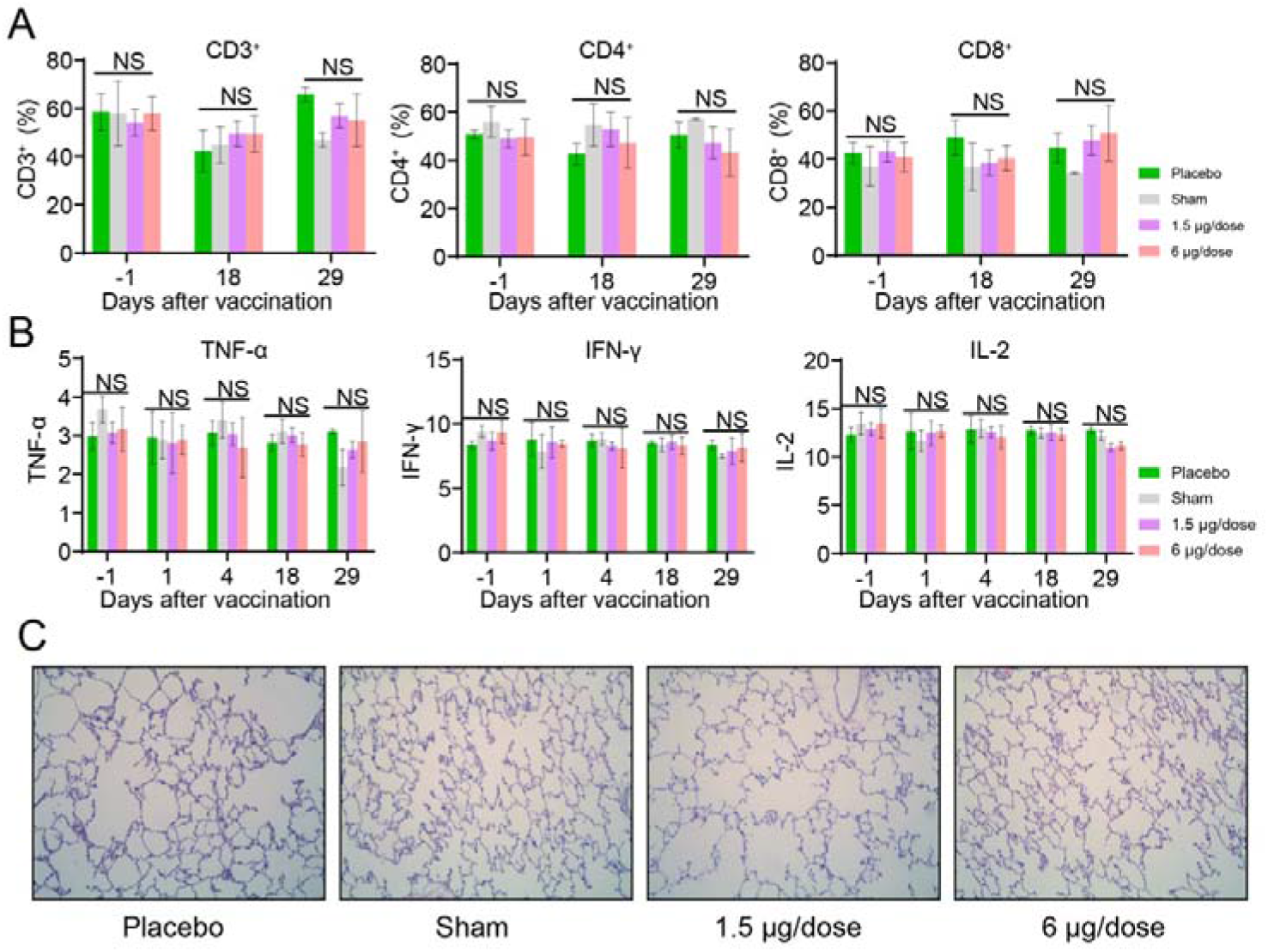
Safety evaluation of PiCoVacc in nonhuman primates. Macaques were immunized three times at day 0, 7 and 14 through the intramuscular route with low dose (1.5 μg per dose) or high dose (6 μg per dose) of PiCoVacc or adjuvant only (sham) or placebo. (**A** and **B**) Hematological analysis in all four groups of macaques (n=4). Lymphocyte subset percents (**A**), including CD3^+^, CD4^+^ and CD8^+^ were monitored at day −1 (1 day before vaccination), 18 (3 day after the second vaccination) and 29 (7 day after the third vaccination). Key cytokines (**B**), containing TNF-α, IFN-γ and IL-2 were examined at day −1, 1 (the day for the first vaccination), 4, 18 and 29 after vaccination. Values are mean ± s.d. (**C**) Histopathological evaluations in lungs from four groups of macaques at day 29. Lung tissue was collected and stained with hematoxylin and eosin.

The serious pandemic of the current COVID 19 and the precipitously increasing numbers of death worldwide necessitate the urgent development of a SARS-CoV-2 vaccine, requiring a new pandemic paradigm. The safety and efficacy are essential for vaccine development at both stages of preclinical studies and clinical trials. Although it’s still too early to define the best animal model for studying SARS-CoV-2 infections, rhesus macaques that mimic COVID-19-like symptoms after SARS-CoV-2 infection appear quite promising animal models for studying the disease. We provide extensive evidences for the safety of PiCoVacc in macaques; neither infection enhancement nor immunopathological exacerbation was observed in our studies. Our data also demonstrate a complete protection against SARS-CoV-2 challenge with a 6μg per dose of PiCoVacc in macaques. Collectively these results suggest a path forward for clinical development of SARS-CoV-2 vaccines for use in humans. Phases I, II and III clinical trials with PiCoVacc, as well as other SARS-CoV-2 vaccine candidates, are expected to begin later this year.

## Acknowledgments

We thank Prof. Zihe Rao and Prof. Junzhi Wang for project discussion, Prof. Weiwei Zhai and Dr. Minhao Li for phylogenetic analysis. We gratefully acknowledge the authors, originating and submitting laboratories of the sequences from GISAID’s EpiFlu™ Database on which this research is based. Work was supported by the National Key Research and Development Program (2020YFC0842100, 2020YFA0707500), the Strategic Priority Research Program (XDB29010000) and the Beijing science and technology plan (Z201100005420006). Xiangxi Wang was supported by Ten Thousand Talent Program and the NSFS Innovative Research Group (No. 81921005).

## Author contributions

L.B., H.M., L.W., K. Xu., Y. L., H.G., X.G., B.K., Y.H., J.L., F.C., D.J., Y.Y., C-F.Q., J.L., X.L., W.S., D.W., H.Z., L.Z., W.D., and Y.L. performed experiments; Q.G., C.Q., Y.Z.,W.Y., X.W., C.L., J-X.L. and X.G. designed the study; all authors analyzed data; and X.W. wrote the manuscript.

## Competing interests

All authors have no competing interests.

## Data and materials availability

The complete genome sequences of SARS-CoV-2 used in this study have been deposited in GenBank under accession number XXX, XXX, XXX, XXX, XXX, XXX, XXX, XXX and XXX, respectively.

## Supplementary Materials for

### Materials and Methods

#### Facility and ethics statements

All experiments with live SARS-CoV-2 viruses were performed in the enhanced biosafety level 3 (P3+) facilities in the Institute of Laboratory Animal Science, Chinese Academy of Medical Sciences (CAMS) approved by the National Health Commission of the People’s Republic of China. All experiments with mice, rats and macaques were carried out in accordance with the Regulations in the Guide for the Care and Use of Laboratory Animals of the Ministry of Science and Technology of the People’s Republic of China.

#### Virus titration

SARS-CoV-2 virus titer was determined by microdose cytopathogenic efficiency (CPE) assay. Serial 10-fold dilutions of virus contained samples were mixed with 1-2 × 10^4^ Vero cells, and then plated in 96-well culture plate. After 3-7 days culture in a 5% CO_2_ incubator at 36.5°C, cells were checked under a microscope for the presence of CPE. Virus titer was calculated by the method of Karber *(20).*

#### Vaccine candidate preparation

SARS-CoV-2 coronavirus, isolated from a COVID-19 infected patient, was provided by Zhejiang Provincial Center for Disease Control and Prevention. Viruses were cultured in large-scale Vero cells factories, and inactivated with β-propiolactone for 24 hours, followed by purification with Ion-exchange Chromatography (IEC) and Size Exclusion Chromatography (SEC) method. The purified viruses were mixed with Al(OH)_3_ adjuvant and served as SARS-CoV-2 vaccine candidate.

#### RT-PCR

Total RNA was extracted from organs as described previously (*13*) with the RNeasy Mini Kit (Qiagen) and the PrimerScript RT Reagent Kit (TaKaRa). The forward and reverse primers targeting against the envelope (E) gene of SARS-CoV-2 used for RT-PCR were 5’-TCGTTTCGGAAGAGACAGGT-3’ and 5’-GCGCAGTAAGGATGGCTAGT-3’, respectively. RT-PCR was performed at the reaction conditions of 50°C for 30 min, followed by 40 cycles of 95°C for 15 min, 94°C for 15 s, and 60°C for 45 s.

#### Vaccine immunogenicity analysis

Balb/c mice, wistar rats were randomly divided into three groups and immunized intraperitoneally and intramuscularly with the trial vaccine at three doses (1.5 μg, 3 μg, 6 μg/dose), respectively. All grouped animals were immunized for two times (at day 0 and 7). The control group was injected with physiological saline. Animals were bled from the tail veins, followed by antibody neutralizing assay to analyze vaccines immunogenicity.

#### Neutralizing assay

Serum samples collected from immunized animals were inactivated at 56°C for 0.5h and serially diluted with cell culture medium in two-fold steps. The diluted serums were mixed with a virus suspension of 100 TCID_50_ in 96-well plates at a ratio of 1:1, followed by 2 hours incubation at 36.5°C in a 5% CO_2_ incubator. 1-2 × 10^4^ Vero cells were then added to the serum-virus mixture, and the plates were incubated for 5 days at 36.5°C in a 5% CO_2_ incubator. Cytopathic effect (CPE) of each well was recorded under microscopes, and the neutralizing titer was calculated by the dilution number of 50% protective condition.

#### Enzyme linked immunosorbant assay (ELISA)

SARS-CoV-2 antibody titer of serum samples collected from immunized animals was determined by indirect ELISA assay. 96-well microtiter plates were coated with 0.1 μg of purified S protein, M protein, N protein individually at 2-8 °C overnight, and blocked with 2% BSA for 1h at room temperature. Diluted sera (1:100) were applied to each well for 2h at 37°C, followed by incubation with goat anti-mouse antibodies conjugated with HRP for 1h at 37°C after 3 times PBS wash. The plate was developed using TMB, following 2M H_2_SO_4_ addition to stop the reaction, and read at 450/630nm by ELISA plate reader for final data.

#### Vaccine safety evaluation

SARS-CoV-2 trial vaccine’s safety was evaluated in macaques. Four groups monkeys (5 female and 5 male monkeys/group) were immunized with high dose (6μg /dose), low dose (1.5μg /dose) vaccine, Al(OH)_3_ adjuvant and physiological saline individually for three times at days 0, 7 and 14. Datasets of many safety related parameters were collected during and after immunization, including clinical observation, body weight, body temperature. Analysis of lymphocyte subset percent (CD3+, CD4+ and CD8+), key cytokines (TNF-α, IFN-γ, IL-2, Il,-2, Il,-4, IL-5_S_ IL-6) and biochemical blood test are also performed in collected blood samples. 60% of monkeys were euthanized at day 18 post immunization, and the left 40% were euthanized at day 29. Organs of lung, heart, spleen, liver, kidney and brain were collected for pathologic analysis.

#### Challenge assay of rhesus macaques

Rhesus macaques (3-4 years old) were divided into four groups and injected intramuscularly with high dose (6 μg/dose), medium dose (3 μg/dose) vaccine, Al(OH)_3_ adjuvant and physiological saline respectively. All grouped animals were immunized at three times (days 0, 7 and 14) before challenged with 10^6^ TCID_50_/ml SARS-CoV-2 virus by intratracheal routes. Macaques were euthanized and lung tissues were collected at 7 days post inoculation (dpi). At day 3, 5, 7 dpi, the throat, and anal swabs were collected. Blood samples were collected on 0, 7, 14, and 21 days post immunization, and 3, 5, 7 dpi for hematological analysis and neutralizing antibody test of SARS-CoV-2. Lung tissues were collected at 7 dpi, and used for RT-PCR assay and histopathological assay.

#### Protein expression and purification

To express the prefusion S ectodomain, a gene encoding residues 1-1208 of COVID-19 S (GenBank: MN908947) with proline substitutions at residues 986 and 987, a “GSAS” substitution at the furin cleavage site (residues 682–685) and a 2×StrepTag was synthesized and cloned into the mammalian expression vector pCAGGS. To express the COVID-19 RBD, residues 319-591 of COVID-19 S were cloned upstream of a 2×StrepTag into the mammalian expression vector pCAGGS. To express the N protein, residues 1-419 (GenBank: QHW06046.1) was cloned into vector pET-28a containing a C-terminal 6×His. S ectodomain and RBD were used to transiently transfect HEK Expi 293F cells (Thermo Fisher) using polyethylenimine. Protein was purified from filtered cell supernatants using StrepTactin resin (IBA) before being subjected to additional purification by size-exclusion chromatography using either a Superose 6 10/300 column (GE Healthcare) or a Superdex 200 10/300 Increase column (GE Healthcare) in 20mM Tris pH 8.0, 200 mM NaCl. The N protein was produced in BL21(DE3) upon the introduction of IPTG. After ultrasonication, the supernatant was loaded over Ni-NTA as manual described (GE Healthcare, Wayne, PA, USA) and eluted with elution buffer (20 mM Tris-HCl, 500 mM NaCl, 200 mM imidazole, pH8.0).

#### Phylogenic tree analysis

Fasta sequences for the COVID-19 were retrieved from the GISAID (https://www.gisaid.org/), NCBI and BIGD (https://bigd.big.ac.cn/ncov) database. After quality control (removing sequences with low quality or short sequence length), 455 sequences were retained for the phylogenetic analysis. By combining 9 target sequences with the sequences from the public databases, we conducted sequence alignment using MAFFT and performed phylogenic reconstruction using IQtree (*21*) with default parameters. The inferred maximum likelihood tree is plotted using ggtree (*22*).

#### Cryo-EM sample preparation

For cryo-EM sample preparation, a 3 μL aliquot of purified viral particles was applied to a glow-discharged C-flat R2/1 Cu grid. Grids were manually blotted for 3 s in 100% relative humidity for plunge-freezing (Vitrobot; FEI) in liquid ethane, as descripted previously *(23).* All samples were examined on a Titan Krios microscope (FEI).

**Fig. S1.**
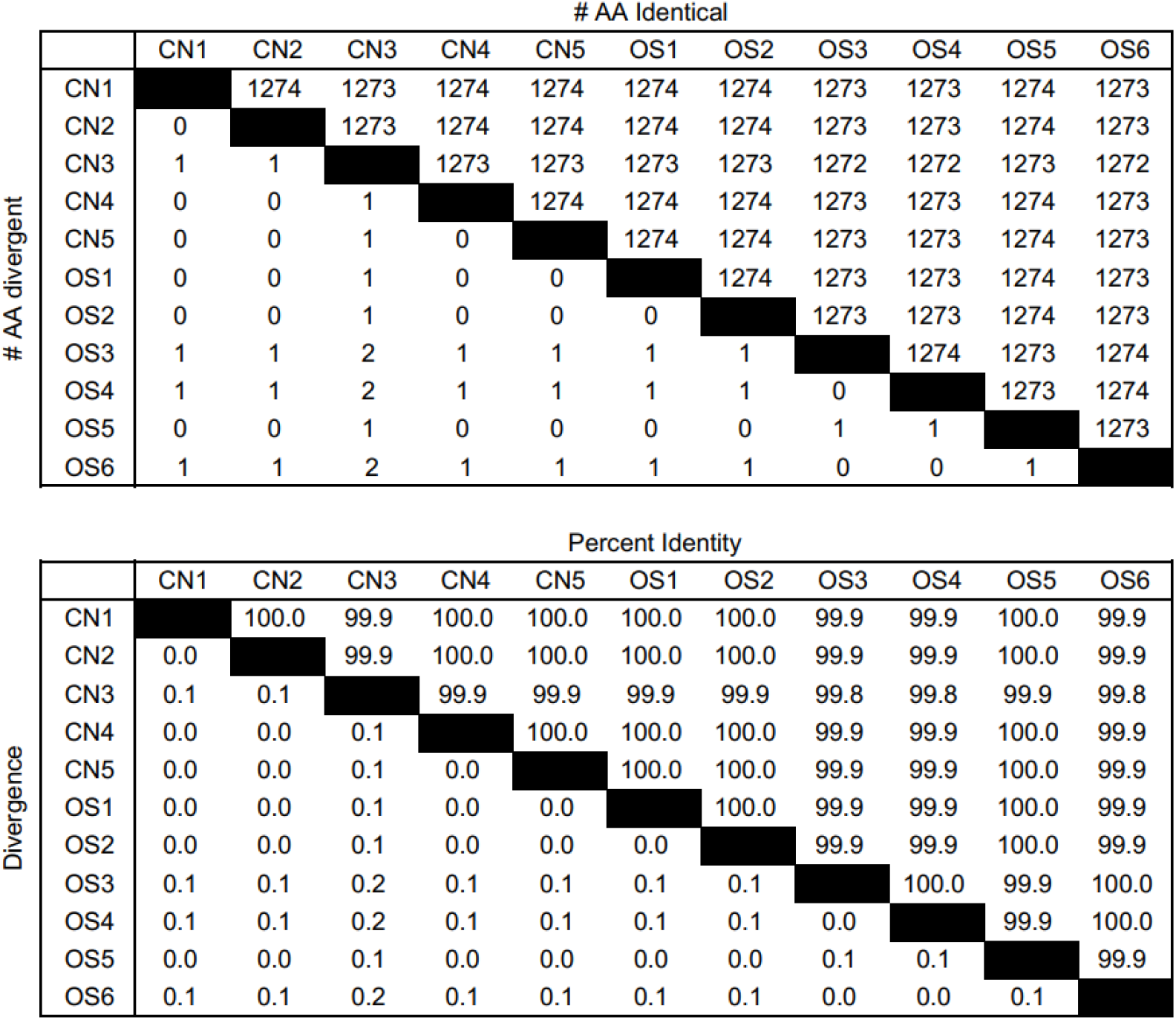
SARS-CoV-2 amino acid sequence comparisons. Number of and percentage of amino acid differences in S are shown for the following SARS-CoV-2 isolates used in this study (Detailed information on these strains is descripted in table S1).

**Fig. S2.**
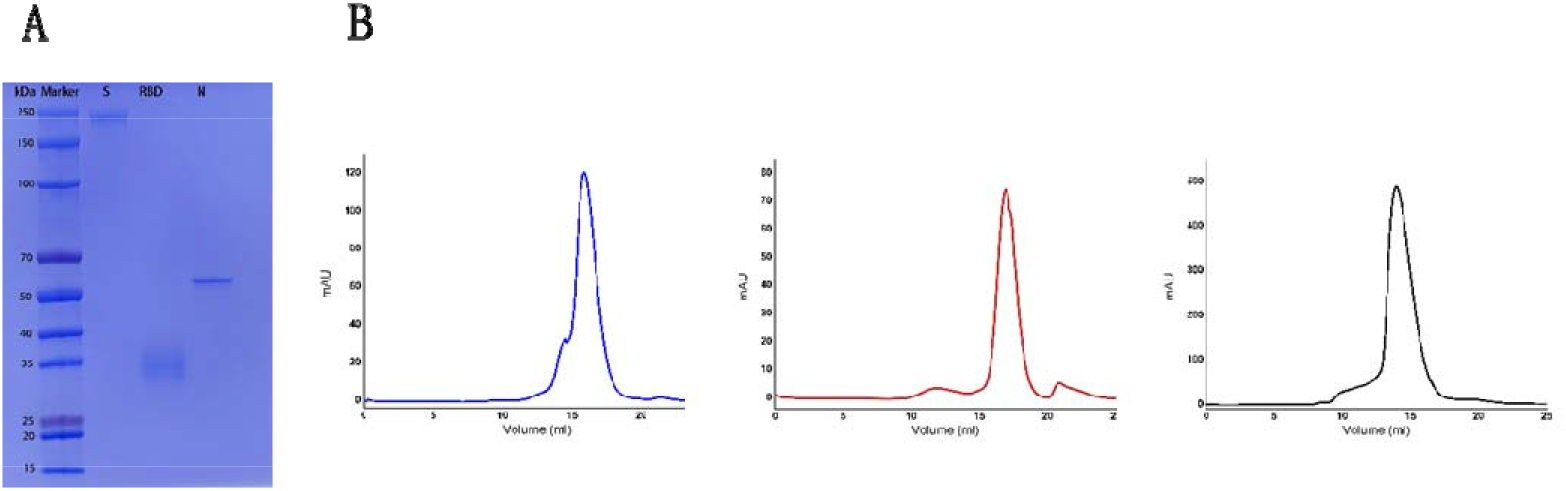
Purification of S, RBD and N protein. **(A)** SDS-PAGE analysis of the S, RBD and N protein. Lane 1: molecular weight ladder, with relevant bands labeled; lane 2: the recombinant S protein; lane 3: the recombinant RBD protein; lane 4: the recombinant N protein. **(B)** Size-exclusion chromatogram of the affinity-purified S, RBD and N protein. (Left) Data of S protein from a Superose 6 10/300 column are shown in red. (Middle) Data of RBD protein from a Superdex 200 10/300 column are shown in red. (Right) Data of N protein from a Superdex 200 10/300 column are shown in black.

**Fig. S3.**
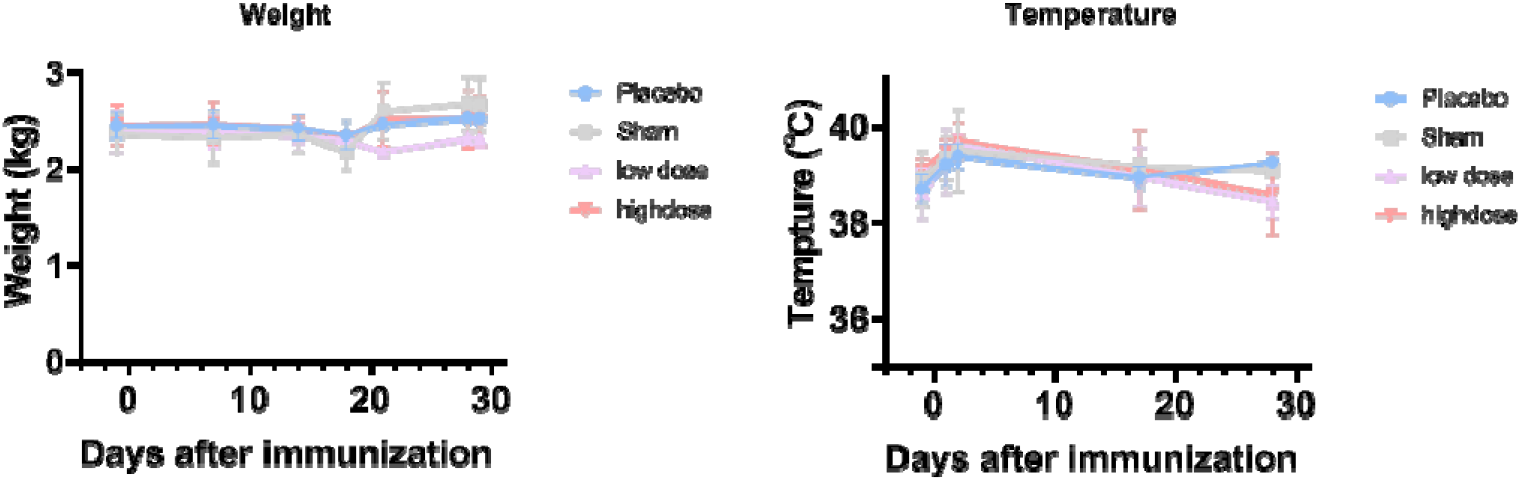
Body weight and body temperature are monitored to evaluate the safety of PiCoVacc in nonhuman primates. Macaques were immunized three times at day 0, 7 and 14 through the intramuscular route with low dose (1.5 μg per dose) or high dose (6 μg per dose) of PiCoVacc or adjuvant only (sham) or placebo. Body weight and body temperature are monitored at different time points.

**Fig. S4.**
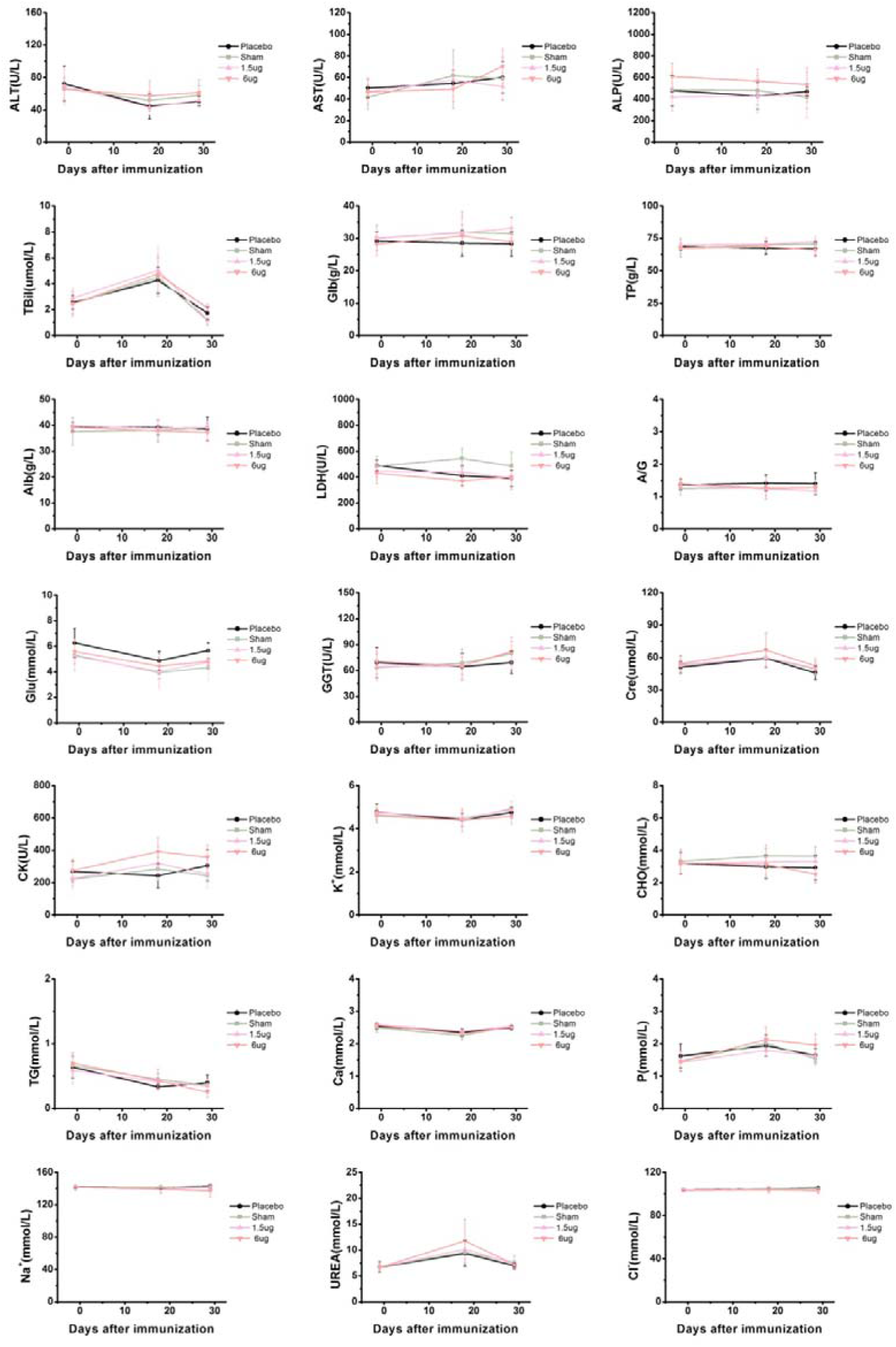
Hematological indices are monitored to evaluate the safety of PiCoVacc in nonhuman primates. Macaques were immunized three times at day 0, 7 and 14 through the intramuscular route with low dose (1.5 μg per dose) or high dose (6 μg per dose) of PiCoVacc or adjuvant only (sham) or placebo. A number of hematological indices are measured at different time points. ALT (Alanine aminotransferase), AST (Aspartate aminotransferase), ALP (Alkaline phosphatase), TBil (Total bilirubin), GGT (γ-glutamyltranspeptidase), TP (Total protein), Alb (Albumin), Glb (Globulin), A/G (Albumin/globulin ratio), UREA (Blood urea), Cre (Creatinine), CK (Creatine kinase), Glu (Glucose), LDH (Lactate dehydrogenase), CHO (Total cholesterol), TG (Triglycerides), Ca (Calcium), P (Phosphorus), Na+ (Sodium ion), K+ (Potassium ion), Cl-(Chloride ion)

**Fig. S5.**
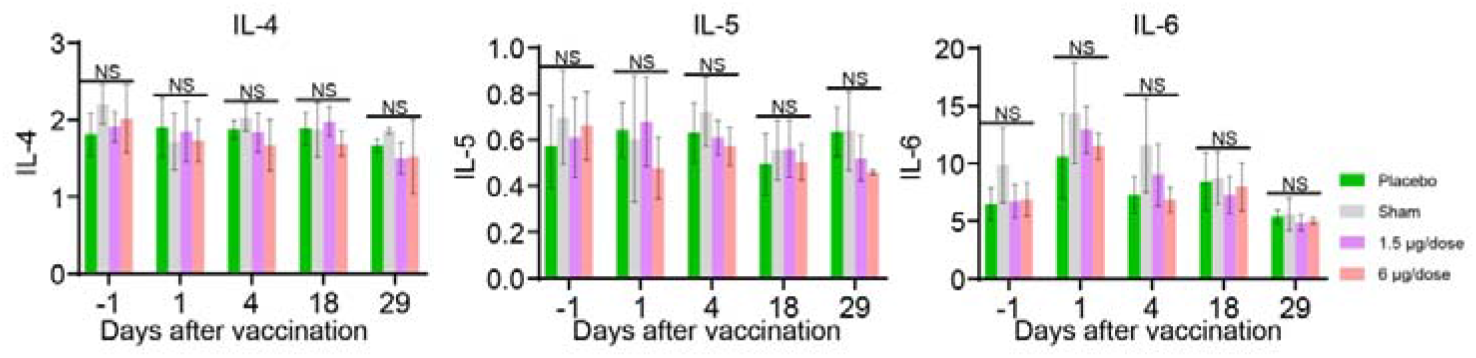
Key cytokines are monitored to evaluate the safety of PiCoVacc in nonhuman primates. Macaques were immunized three times at day 0, 7 and 14 through the intramuscular route with low dose (1.5 μg per dose) or high dose (6 μg per dose) of PiCoVacc or adjuvant only (sham) or placebo. IL4, IL5 and IL6 are measured at different time points.

**Fig. S6.**
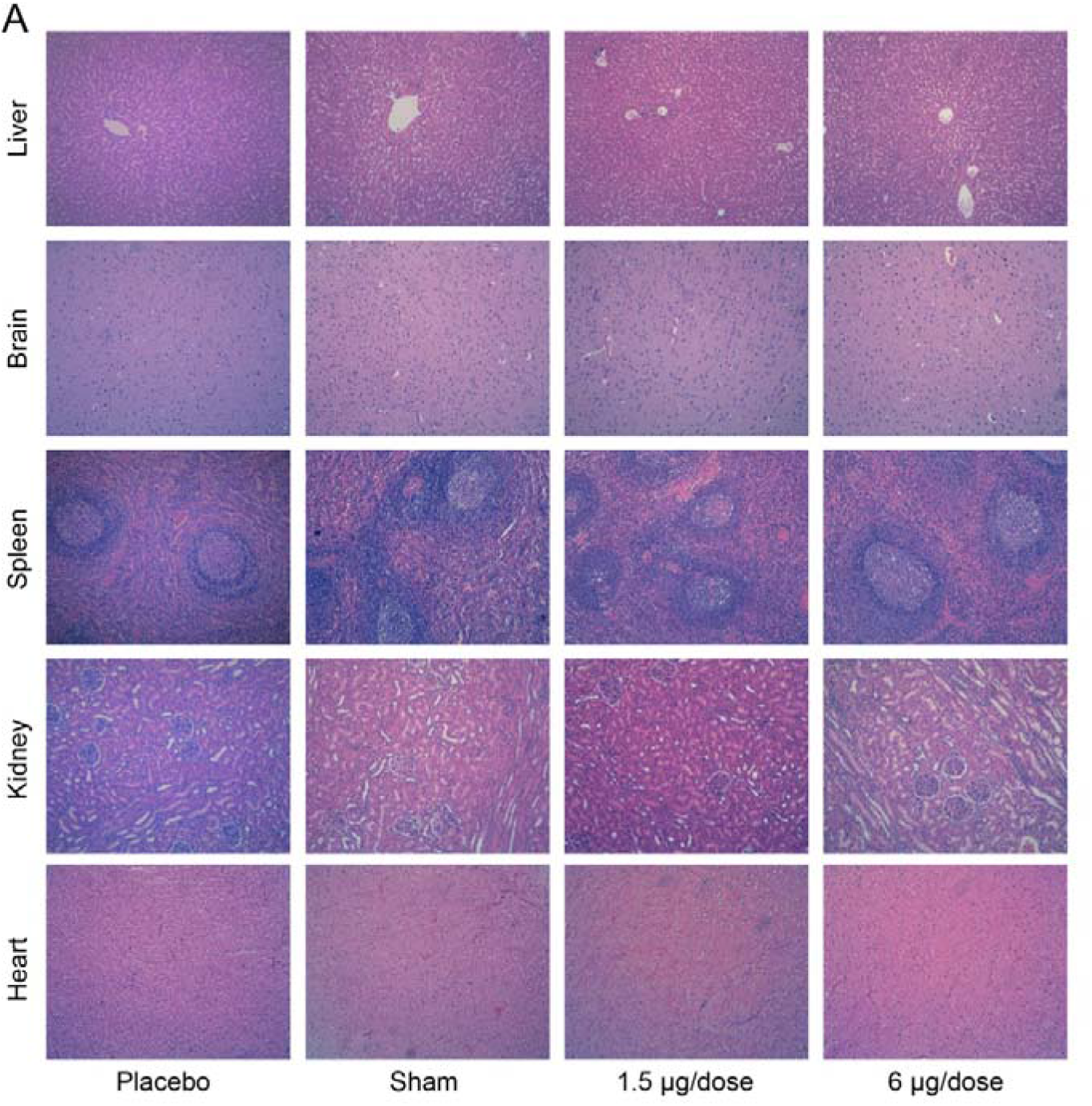
Histopathological evaluations are performed to evaluate the safety of PiCoVacc in nonhuman primates. Macaques were immunized three times at day 0, 7 and 14 through the intramuscular route with low dose (1.5 μg per dose) or high dose (6 μg per dose) of PiCoVacc or adjuvant only (sham) or placebo. Histopathological evaluations in brain, spleen, kidney and heart from four groups of macaques at day 29. Tissues were collected and stained with hematoxylin and eosin.

**Table S1.** Virus strain information studied in this manuscript

| Strain | Country where virus isolated | GenBank accession number |
| --- | --- | --- |
| CN1 | China | xxx |
| CN2 | China | xxx |
| CN3 | China | xxx |
| CN4 | China | xxx |
| CN5 | China | xxx |
| OS1 | Italy | xxx |
| OS2 | Italy | xxx |
| OS3 | Spain | xxx |
| OS4 | Switzerland | xxx |
| OS5 | United Kingdom | xxx |
| OS6 | Italy | xxx |

**Table S2.** Genetic stability analysis of PiCoVacc. The sequence initially determined for PiCoVacc P1 stock was used as a reference sequence.

| Stock | Mutations identified in structural proteins |  | Additional mutations identified in genome |  |
| --- | --- | --- | --- | --- |
|  | genome nt position | amino acid | genome nt position | amino acid |
| P1 | -- | -- | -- | -- |
| P3 | C26309A | E-A32D | none | none |
| P5 | C26309A | E-A32D | C13170T | nsp10-T49I |
| P10 | C26309A | E-A32D | C13170T | nsp10-T49I |

## Notes

### Competing Interest Statement

The authors have declared no competing interest.

